# Revealing the hidden social structure of pigs with AI assisted automated monitoring data and social network analysis

**DOI:** 10.1101/2025.01.30.635669

**Authors:** Saif Agha, Eric Psota, Simon P. Turner, Craig R. G. Lewis, Juan P Steibel, Andrea Doeschl-Wilson

**Affiliations:** The Roslin Institute, University of Edinburgh, Easter Bush, Edinburgh EH25 9RG, UK; Animal Production Department, Faculty of Agriculture, Ain Shams University, Shubra Alkhaima, Cairo 11241, Egypt; PIC North America, Hendersonville, TN, USA; Animal and Veterinary Sciences Department, Scotland’s Rural College, West Mains Road, Edinburgh EH9JG, UK; PIC, C/Pau Vila no. 22, Sant Cugat del Valles, 08174 Barcelona, Spain; Department of Animal Science, Iowa State University, Ames, IA 50011, USA

**Keywords:** Deep learning, Automated monitoring, Digital phenotypes, Social networks, farm animals

## Abstract

Social interactions of farm animals affect their performance, health and welfare. The recent advances in AI-automated monitoring technologies offer digital phenotypes, at low-cost, that record the animals in real-time. This proof-of-concept study addresses, for the first time, the hypothesis that applying social network analysis (SNA) on automated data could potentially facilitate the analysis of social structures of farm animals. Data was collected using automated recording systems that captured 2D camera images and videos of pigs in six pens (16–19 animals each) on a PIC breeding company farm (USA). The system provided real-time data, including ear-tag readings, elapsed time, posture (standing, lying, sitting), and XY coordinates of the shoulder and rump for each pig. Weighted SNA was performed, based on the proximity of “standing” animals, for two 3-day periods: the early growing period (first month after mixing) and the later period (60 days post-mixing). Group level degree, betweenness, and closeness centralization showed a significant increase from the early growing period to the later one (p<0.02), highlighting the pigś social dynamics over time. Largest clique size remained unchanged (p=0.28), but the number of maximal cliques significantly decreased from the early to late growing period (p=0.007). Individual SNA traits were stable over these periods, except for closeness centrality and clustering coefficient which significantly increased (p<0.00001). This study demonstrates that combining AI-assisted monitoring technologies with SNA offers an efficient, real-time approach to gain novel insights into animal social interactions. This approach can optimize on-farm management or breeding practices, leading to improved animal performance, health, and welfare.

**Highlights:**

- Social network analysis (SNA) applied to automated monitoring data provides novel insight into social interactions of pigs.
- Group level SNA centralization traits highlighted pigs ‘social dynamics over time.
- The stability of key individual SNA traits can be leveraged for breeding purposes.
- AI-assisted monitoring combined with SNA help optimizing management practices on commercial farms.

## 1. Introduction

Livestock and aquaculture species are usually housed in groups on breeding and commercial farms. Within these groups, both positive and negative social interactions play a significant role in shaping animal behaviour, welfare, productivity, and health. Specifically, in pigs, negative social interactions have been found to impact access to resources, feeding behaviour, and overall growth [1,2]. Moreover, these interactions can negatively affect welfare and stress levels, particularly when harmful behaviours, such as aggression or tail biting, occur [3,4]. Furthermore, both socially positive and negative forms of interaction, which involve close proximity, usually increase the risk of disease transmission [4,5]. Therefore, quantifying the structure and dynamics of social interactions within groups could provide crucial information to improve animal productivity, health, and welfare [6,7].

In this context, social network analysis (SNA) has become an essential tool for investigating the complex social structures of animal groups [8]. It enables the quantification of group-level social measures, e.g. centralization, which provide insights into the social structure of animal groups [9,10]. This method also facilitates the identification of subgroups, e.g., communities and cliques (representing fully connected individuals) [11], that share specific attributes and behaviour [12]. SNA also provides individual-level traits that quantify the role of each animal within its social network [13,14]. This establishes SNA as a comprehensive tool for understanding social structures and their dynamics within commercial animal populations [15].

In pigs, SNA applied to manually extracted data from video recordings of group-housed individuals, has revealed both the direct and indirect roles of each animal in pen-level aggression and association with resulting skin lesions[16–18]. Furthermore, research has demonstrated that SNA traits, quantifying individual centrality in aggression networks, are heritable and have favourable correlations, both at the phenotypic and genetic level, with economically important traits [16,19]. These findings suggest that SNA could help in the development of advanced breeding strategies to simultaneously improve animal performance and welfare. In practice, however, manual decoding of video images to study animal social interactions is time-consuming and labour-intensive and is prone to observer bias [20]. Recently, automated monitoring systems integrated with advanced artificial intelligence technologies, such as deep learning (DL), have shown a considerable promise in improving the efficiency of observation and detection of farm animals and their behaviour [21]. The application of DL and machine vision approaches in pigs has been shown to have high accuracy under farm conditions in identifying postures such as standing, sitting, and lying, as well as daily activities, like eating and drinking [22]. Furthermore, tracking technologies have offered digital phenotypes, at low-cost, that record the movement and coordinates of the animals in real-time under commercial farm conditions [23,24]. Compared to traditional animal monitoring methods, DL methods do not need to manually extract features but take advantage of convolutional neural networks (CNN) to replace the traditional image processing, achieve a high accuracy in monitoring animals under different farm conditions [25].

This proof-of-concept study addresses, for the first time, the hypothesis that applying SNA approaches on automated data could potentially facilitate the analysis of social structures of animal populations and gain insights into their social interactions. The objectives of this study were to (1) evaluate the feasibility of using data from an on-farm computer monitoring system in commercial pigs coupled with DL to construct social networks, (2) gain new insights into the social structure and its dynamic changes within pens, and (3) identify the role of each animal in the social interactions occurring in groups of growing pigs in commercial farms.

## 2. Materials and Methods

### 2.1. Automated data

Data was derived from automated recording systems that provide 2D camera images (See Figure 1) and video recordings of pigs in six pens, each containing 16-19 purebred animals, at a nucleus farm of the PIC breeding company in the USA. The automated system provides real-time data from reading the ear tag, the number of seconds that has elapsed since the start of video recording, as well as the position and posture e.g., standing, lying and sitting for each animal. Video footage was processed using a multi-object tracking algorithm to extract individualized position and activity data [24]. Initially, a customized version of the DeepCut pose estimation algorithm detected key points on the pigs, such as midpoints, snouts, and right ear tag locations, and linked them to individual animals [26]. A CNN then identified each pig’s posture and activities (e.g., eating or drinking). Each pig was tracked frame-by-frame using the Hungarian algorithm [27], with any undetected pigs assumed to remain stationary. Reliable tracking was further supported by a custom ear-tag reading method developed by PIC [28,29]. The automated system also utilized DL algorithms to generate 2D XY coordinates for the front (shoulder) and rear (rump) positions of each pig, with measurements normalized to meters from the top-left corner of each pen (Figure 1). To ensure accurate data collection, a nine-month “learning period” was established for equipment fine-tuning and procedure optimization. Data from this period were excluded from analysis to prevent bias and ensure recording consistency (See details in [24]). Details on the system’s validation for accurately identifying individual animals’ postures (lying, sitting, standing), activities (eating, drinking), and positions (XY coordinates of the shoulder and rump) are provided in the supplementary material.

**Figure 1.**
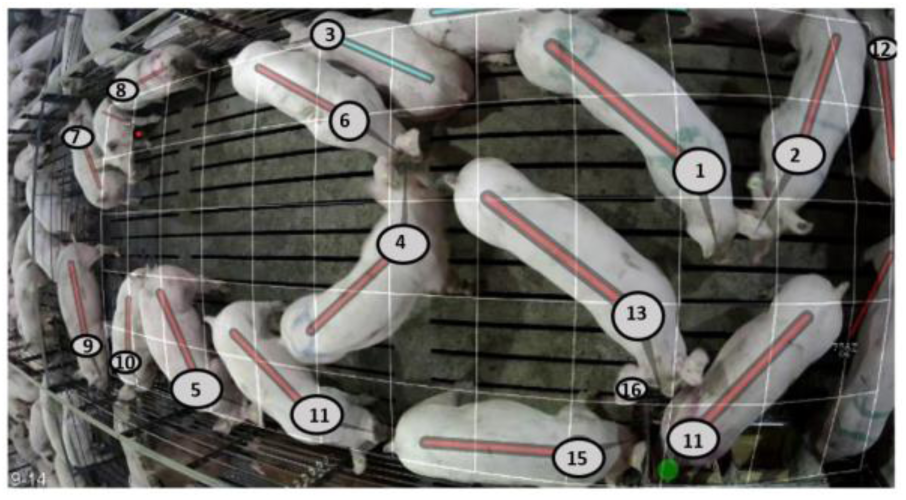
The 2D camera images provided by the automated recording systems which shows the ear tag, position and posture, e.g., standing (with red line) and sitting (blue line) for each pig within the pen (Animalś ID have been anonymised for this illustration).

### 2.2. SNA

Social networks were constructed based on the automated tracking data to describe the contact structure of each of the six pens. To construct these networks and for subsequent SNA, a computational pipeline was developed using various R packages, i.e., *spatsoc* [30], *asnipe* [31], *sna* [32], *vegan* [33], and *igraph* [34]. In this study, only “standing” animals were included in the social network construction. This approach enabled the investigation of social interactions only during active behaviour, providing insights into the social dynamics within a pen. For each pen, a total of 12 hours per day of continuous interaction time between individuals was included in the SNA, spanning from 06:00 to 18:00, over three consecutive days. Furthermore, SNA was conducted for two separate 3-day periods; the early growing period (the first month after mixing) and a later growing period (60 days after mixing) to allow for examination of the temporal changes in the social structure within the growing period. The average “standing” time/day for the pigs was 170.28 minutes (SD= 52.21 minutes)[24]. The XY coordinates of the pigs’ shoulders, along with timestamps provided by the automated system, were used to define the interactions of pigs within contemporary groups spanning the time period of 12 hours per day for social network construction. Subsequently, a spatial analysis was performed to identify animals that interactedbased on their proximity within this contemporary group. Here, proximity between animals within the same contemporary group was defined by a threshold of 0.5 meters Euclidean distance between the shoulders, sustained for a duration longer than average interaction time between individuals within each pen each day (Table S2). The chosen threshold of 0.5 meters accommodates for potential measurement errors in the XY coordinates (see validation) and is within the range of previously identified proximity distances between 0.3 to 1 meter defining interactions [35]. Based on the proximity of dyadic interactions, an undirected social network was constructed for each day (i.e. three consecutive days for both the early and late growing period for each pen). These networks were weighted based on the duration of proximity interactions between each dyad. Group-level SNA traits, e.g., group-degree centralization, group-closeness centralization, group-eigenvector centralization, group-betweenness centralization, density and modularity, were then computed, and combined over the three days from the social networks to quantify the social structure of each pen for each of the two growing periods. Temporal differences in these group-level SNA traits were then assessed using linear mixed models, with pen and day of measurement as random effects, growing period (early/ late) as a fixed effect and pen size as a covariate. To further investigate the social dynamics and gain detailed insights into the social structure of the pen, community detection and clique analyses were performed for the social networks. For community detection, one of the most common methods used in animal populations was applied i.e., the modularity-based methods implementing the Louvain algorithm [36]. For the clique analysis, the number of maximal cliques and the size of the largest clique were computed. Then, the temporal differences in the number of communities, the number of maximal cliques and the size of the largest clique were assessed using the same linear mixed models as described above. Due to the importance of cliques in the social structure, further analysis of the maximal cliques was also performed by identifying the co-membership matrix of these cliques [37]. The Mantel test was then performed to compare the clique co-membership matrices between days, and between the early and later growing periods, to provide insight into the development of cliques among pigs over the growth stages [33]. Finally, individual SNA traits were computed for each individual per day for the different growing periods. The definitions of the terminology and diverse SNA traits used in this study are shown in Table 1.

**Table 1.** Definition of the terminology and diverse SNA traits used in this study.

| Measure | Definition |
| --- | --- |
| Proximity | Proximity was defined by a threshold of 0.5 meters Euclidean distance between the shoulders of the “standing” pigs, which was sustained for a duration longer than the average interaction time between each pair of individuals within each pen each day. |
| Weighted SNA graph | A representation of a network where the edges (interactions) between nodes (animals) are assigned numerical values (weights) that reflect the duration of the interactions (proximity). |
| Individual degree centrality | The number of edges (i.e. interactions) attached to a node (animal). |
| Individual betweenness centrality | Betweenness centrality in a weighted network measures how often a node (or individual) acts as a bridge along the shortest paths between other nodes. |
| Individual closeness centrality | The sum of the direct connections between a focal node and other nodes in the network. |
| Individual eigenvector centrality | The connectivity of a node according to the all-degree centrality of the node and the all-degree centrality of the nodes that it connects with <sup>[38]</sup>. |
| Individual clustering coefficient | The proportion of an individual node's connections that are also connected with each other relative to the number of theoretically possible connections <sup>[10]</sup>. |
| Centralization | A graph level centralization is computed from the individual centrality scores of the nodes using the formula,<br>$\text{Centralization} = \frac{\sum_{i=1}^n (C_{\max} - C_i)}{\text{maximum possible centralization}}$<br>where Cmax is the centrality of the most central node. Ci is the centrality of node, and the centralization was normalized by dividing by the maximum theoretical score for a graph with the same number of nodes <sup>[39]</sup>. For degree, closeness and betweenness centralization the most centralized structure is a network where a small number of nodes hold most of the connections. For eigenvector centralization, a high centralization graph indicates that a small number of nodes have a very high eigenvector centrality, while the majority have much lower values. |
| Density | The ratio of the number of edges present in the network relative to the total number of possible edges in a group of the same size. |
| Modularity | The strength of the division of a network into communities. It evaluates how well the network is divided into subgroups, where nodes within the same subgroup (or community) are more densely connected to each other than to nodes in other groups <sup>[40]</sup>. |
| Community detection | The process of identifying subgroups or clusters of nodes within a network that are more densely connected internally compared to the connections in the network <sup>[40]</sup>. |
| Modularity-based community detections method | The modularity-based method, using Louvain Algorithm for community detections, aims to maximize the modularity score to identify densely connected subgroups within the network <sup>[41]</sup>. |
| Clique | A subset of nodes where every node is directly connected to every other node in the subset. |
| Maximal clique | A clique is maximal if there is no node in the graph that can be added to this clique to create a larger clique without violating the clique property <sup>[42]</sup> |
| Co-membership | The relationship between nodes based on shared membership in specific cliques. |
| Largest clique size | A clique with the maximum number of nodes among all cliques in that graph <sup>[42]</sup>. |

## 3. Results

The above-described SNA pipeline applied to the automated data produced weighted networks based on the proximity between animals in each pen within the two growth periods. Figure 2 shows SNA graphs for one of six pens, illustrating the connections reflecting pairwise interactions weighted by duration, the communities identified by the modularity-based method, members of the largest clique, and co-membership density for the maximal cliques.

**Figure 2:**
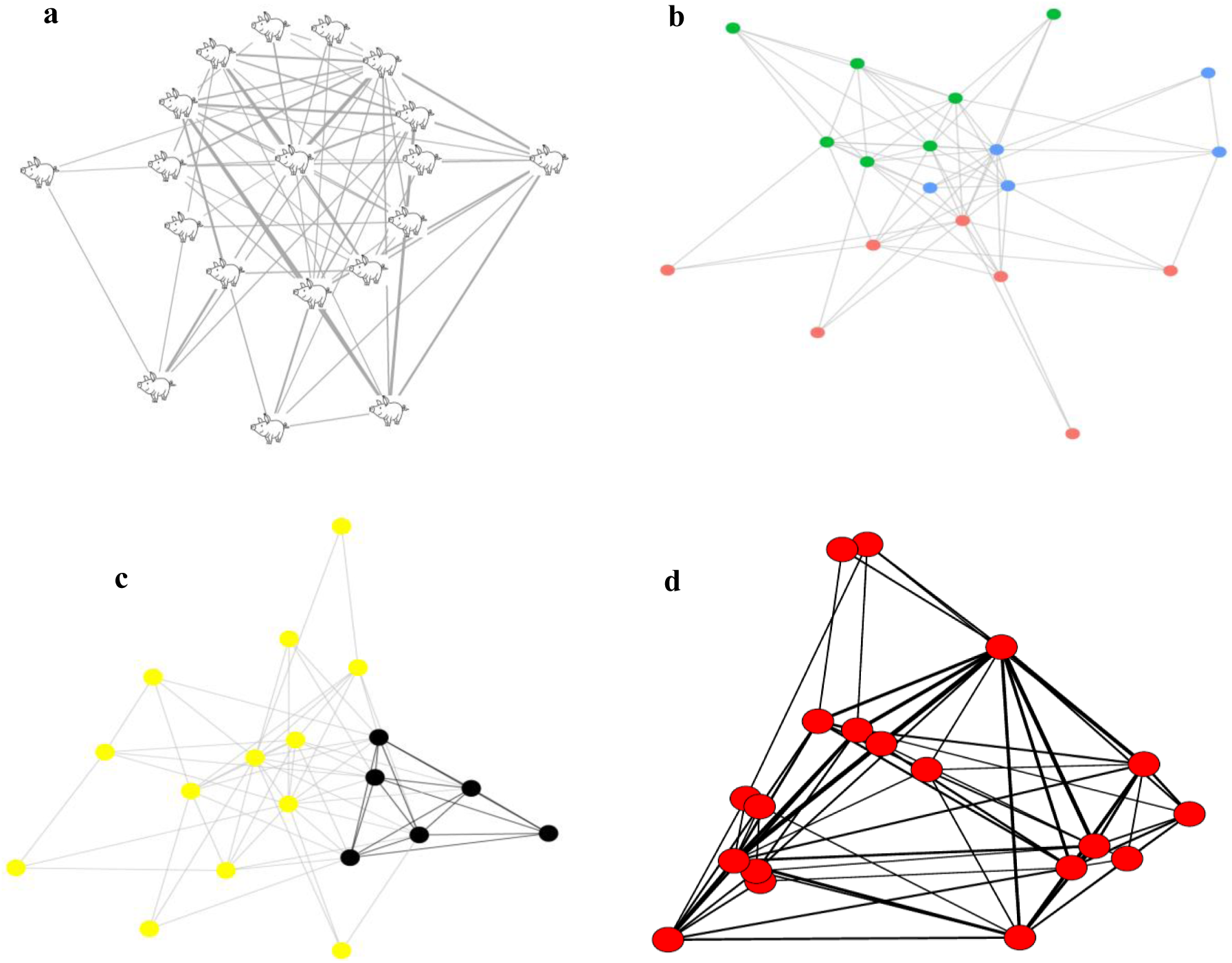
Social network graphs for one of the studied pens for one of the studied days, showing: a) a network where connections (edges) represent proximity between nodes (animals), with the thickness of the edges indicating the duration of pairwise proximity; b) the same network analysed using a modularity-based method to identify distinct communities, represented by coloured nodes, within the network; c) the same network highlighting key animals, shown in black, that are members of the largest clique, and d) the same network analysed for co-membership in maximal cliques illustrating high-density areas, associated with thick edges, between individuals that frequently co-occurred in maximal cliques.

### 3.1. Group-level SNA traits

The group-level SNA traits for the early and later growing periods of the studied pens are shown in Table 2. The least square means showed a significant difference in these group-level SNA traits between the two growing periods (Figure 3). For instance, group level degree, betweenness, and closeness centralization showed a significant increase from the early growing period to the later growing period (p<0.02). That would indicate that in the later growing period a few individuals had very high centrality measures, while the majority had much lower values. In contrast, a non-significant increase was found for eigenvector centralization (p = 0.12). Also, the density of the network remained stable across both periods (p = 0.79), suggesting no significant change in the overall frequency and duration of social interactions between the pigs in different growing periods.

**Figure 3:**
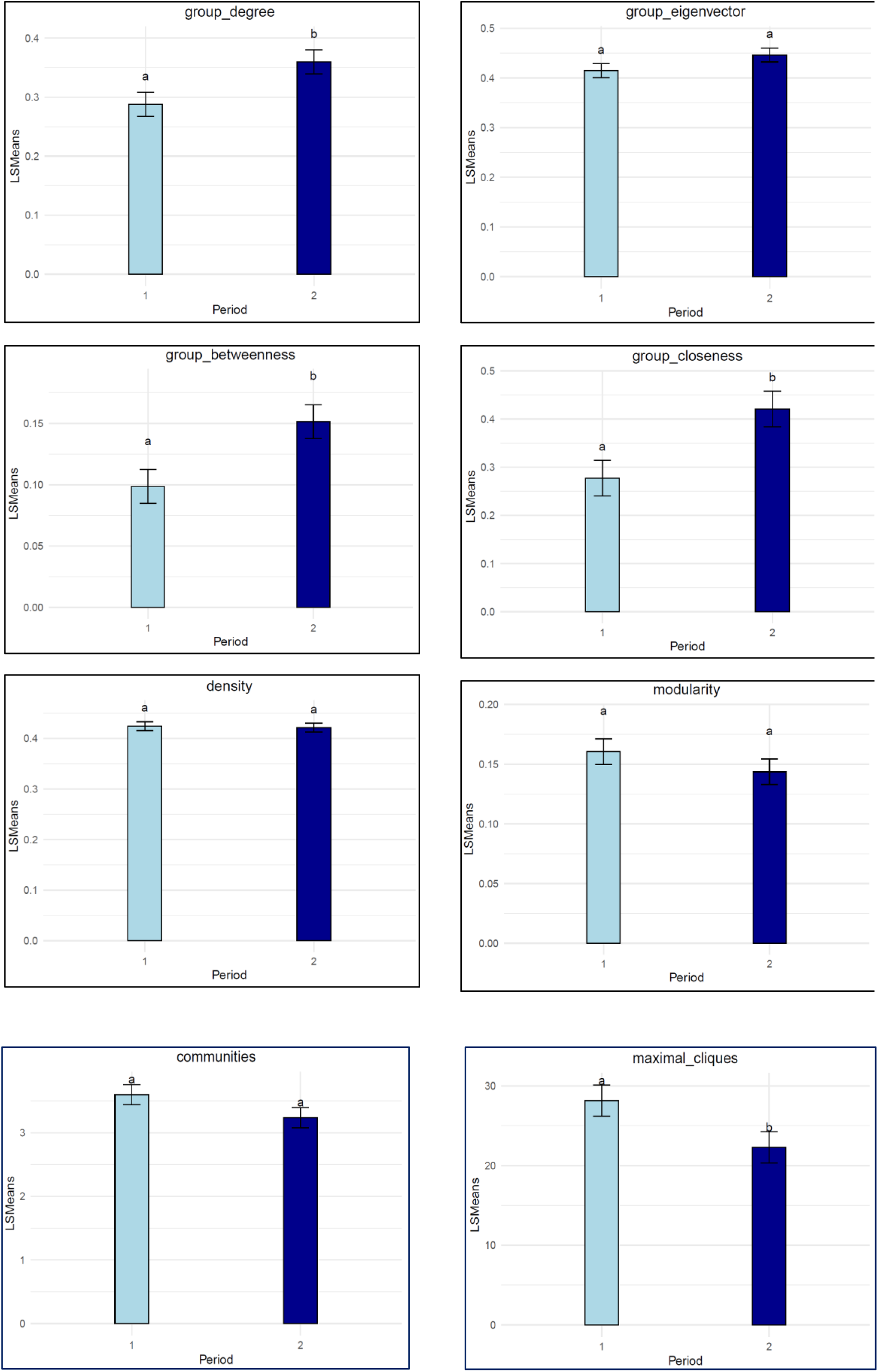
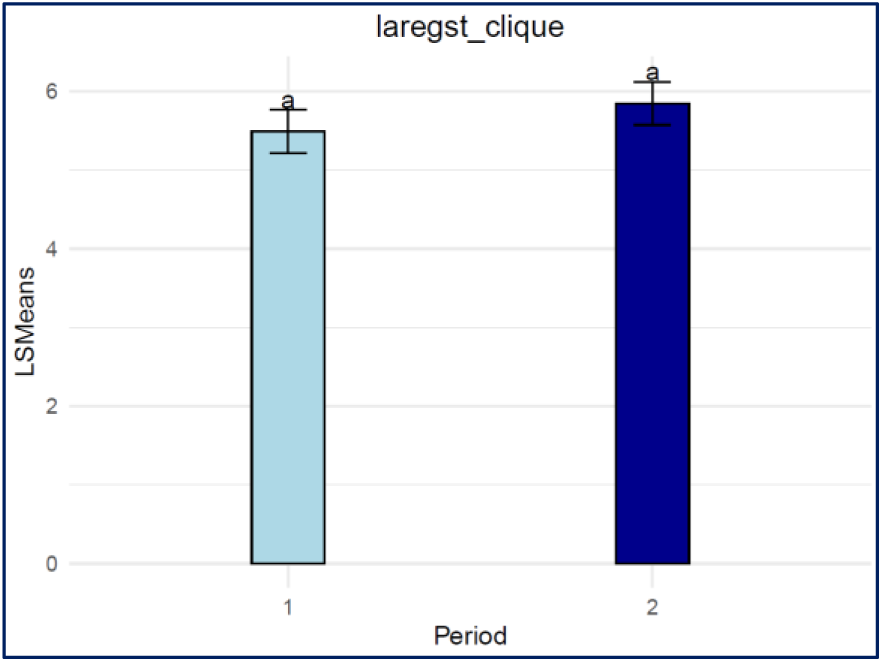
Least square means of group level SNA traits, communities, and clique traits by growing period. Growing periods not connected by the same letter are significantly different at (p < 0.05).

**Table 2:** Descriptive statistics of the group-level SNA traits over 3-days each for the early and later growing periods.

| Pen | Period | Group degree |  |  | Group closeness |  |  | Group eigenvector |  |  | Group betweenness |  |  | Density |  |  | Modularity |  |  |
| --- | --- | --- | --- | --- | --- | --- | --- | --- | --- | --- | --- | --- | --- | --- | --- | --- | --- | --- | --- |
|  |  | Mean (SD) | Max | Min | Mean (SD) | Max | Min | Mean (SD) | Max | Min | Mean (SD) | Max | Min | Mean (SD) | Max | Min | Mean (SD) | Max | Min |
| 1 | 1 | 0.36<br>(0.08) | 0.44 | 0.28 | 0.30<br>(0.27) | 0.50 | 0.00 | 0.48<br>(0.06) | 0.53 | 0.41 | 0.12<br>(0.07) | 0.18 | 0.04 | 0.40<br>(0.04) | 0.44 | 0.36 | 0.15<br>(0.06) | 0.22 | 0.10 |
| 1 | 2 | 0.37<br>(0.08) | 0.47 | 0.32 | 0.42<br>(0.12) | 0.56 | 0.35 | 0.45<br>(0.01) | 0.45 | 0.45 | 0.18<br>(0.05) | 0.24 | 0.14 | 0.40<br>(0.02) | 0.42 | 0.39 | 0.12<br>(0.04) | 0.15 | 0.08 |
| 2 | 1 | 0.25<br>(0.06) | 0.32 | 0.20 | 0.29<br>(0.09) | 0.39 | 0.23 | 0.37<br>(0.05) | 0.42 | 0.33 | 0.09<br>(0.01) | 0.10 | 0.08 | 0.44<br>(0.02) | 0.46 | 0.42 | 0.19<br>(0.02) | 0.20 | 0.17 |
| 2 | 2 | 0.3<br>(0.13) | 0.45 | 0.22 | 0.35<br>(0.18) | 0.55 | 0.23 | 0.42<br>(0.04) | 0.46 | 0.39 | 0.14<br>(0.1) | 0.25 | 0.08 | 0.42<br>(0.01) | 0.43 | 0.41 | 0.14<br>(0.03) | 0.17 | 0.11 |
| 3 | 1 | 0.31<br>(0.1) | 0.41 | 0.22 | 0.34<br>(0.12) | 0.48 | 0.24 | 0.40<br>(0.07) | 0.47 | 0.33 | 0.12<br>(0.05) | 0.18 | 0.08 | 0.42<br>(0.01) | 0.42 | 0.42 | 0.15<br>(0.03) | 0.18 | 0.12 |
| 3 | 2 | 0.26<br>(0.09) | 0.36 | 0.20 | 0.32<br>(0.11) | 0.45 | 0.24 | 0.43<br>(0.04) | 0.46 | 0.39 | 0.11<br>(0.04) | 0.15 | 0.08 | 0.41<br>(0.03) | 0.44 | 0.38 | 0.18<br>(0.02) | 0.19 | 0.16 |
| 4 | 1 | 0.31<br>(0.03) | 0.34 | 0.28 | 0.13<br>(0.22) | 0.38 | 0.00 | 0.41<br>(0.02) | 0.43 | 0.40 | 0.10<br>(0.06) | 0.16 | 0.04 | 0.44<br>(0.01) | 0.44 | 0.43 | 0.13<br>(0.04) | 0.17 | 0.10 |
| 4 | 2 | 0.37<br>(0.04) | 0.40 | 0.33 | 0.45<br>(0.02) | 0.46 | 0.43 | 0.45<br>(0.03) | 0.47 | 0.41 | 0.16<br>(0.03) | 0.18 | 0.12 | 0.44<br>(0.04) | 0.48 | 0.41 | 0.14<br>(0.03) | 0.17 | 0.12 |
| 5 | 1 | 0.27<br>(0.01) | 0.28 | 0.26 | 0.30<br>(0.03) | 0.32 | 0.27 | 0.42<br>(0.03) | 0.46 | 0.40 | 0.09<br>(0.02) | 0.10 | 0.07 | 0.41<br>(0.03) | 0.44 | 0.39 | 0.18<br>(0.01) | 0.18 | 0.17 |
| 5 | 2 | 0.39<br>(0.01) | 0.40 | 0.38 | 0.45<br>(0.01) | 0.46 | 0.44 | 0.47<br>(0.05) | 0.51 | 0.42 | 0.14<br>(0.04) | 0.18 | 0.10 | 0.41<br>(0.02) | 0.43 | 0.39 | 0.15<br>(0.03) | 0.17 | 0.12 |
| 6 | 1 | 0.33<br>(0.08) | 0.42 | 0.26 | 0.37<br>(0.11) | 0.49 | 0.29 | 0.43<br>(0.07) | 0.50 | 0.36 | 0.10<br>(0.05) | 0.15 | 0.06 | 0.44<br>(0.02) | 0.46 | 0.42 | 0.14<br>(0.02) | 0.15 | 0.12 |
| 6 | 2 | 0.38<br>(0.04) | 0.42 | 0.34 | 0.47<br>(0.06) | 0.54 | 0.42 | 0.45<br>(0.01) | 0.45 | 0.44 | 0.16<br>(0.03) | 0.18 | 0.12 | 0.44<br>(0.02) | 0.46 | 0.42 | 0.16<br>(0.06) | 0.22 | 0.11 |

### 3.2. Communities and cliques

The descriptive statistics of the number of communities, number of maximal cliques, and largest clique size are shown in Table 3. No significant difference was observed for the mean number of communities between the early and late growing period (p = 0.14) (Figure 3). Similarly, there was no change in the largest clique size from the early to the late growing period (p = 0.28). However, there was a significant decrease in the number of maximal cliques between the early and late growing periods (p = 0.007).

**Table 3:**
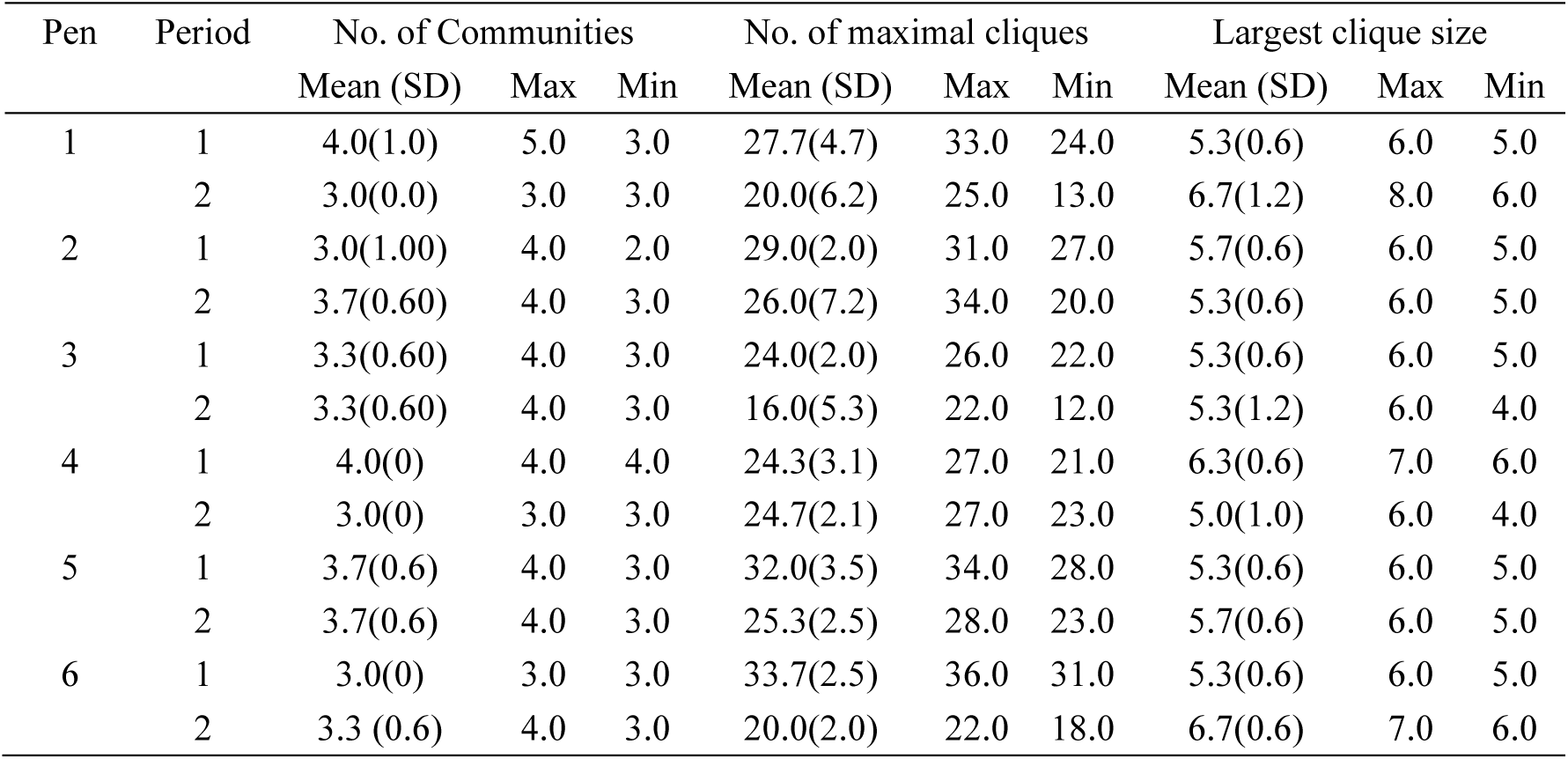
Descriptive statistics of the number of communities, number of maximal cliques, and largest clique size over the 3-days for the early and late growing periods.

| Pen | Period | No. of Communities |  |  | No. of maximal cliques |  |  | Largest clique size |  |  |
| --- | --- | --- | --- | --- | --- | --- | --- | --- | --- | --- |
|  |  | Mean (SD) | Max | Min | Mean (SD) | Max | Min | Mean (SD) | Max | Min |
| 1 | 1 | 4.0(1.0) | 5.0 | 3.0 | 27.7(4.7) | 33.0 | 24.0 | 5.3(0.6) | 6.0 | 5.0 |
|  | 2 | 3.0(0.0) | 3.0 | 3.0 | 20.0(6.2) | 25.0 | 13.0 | 6.7(1.2) | 8.0 | 6.0 |
| 2 | 1 | 3.0(1.00) | 4.0 | 2.0 | 29.0(2.0) | 31.0 | 27.0 | 5.7(0.6) | 6.0 | 5.0 |
|  | 2 | 3.7(0.60) | 4.0 | 3.0 | 26.0(7.2) | 34.0 | 20.0 | 5.3(0.6) | 6.0 | 5.0 |
| 3 | 1 | 3.3(0.60) | 4.0 | 3.0 | 24.0(2.0) | 26.0 | 22.0 | 5.3(0.6) | 6.0 | 5.0 |
|  | 2 | 3.3(0.60) | 4.0 | 3.0 | 16.0(5.3) | 22.0 | 12.0 | 5.3(1.2) | 6.0 | 4.0 |
| 4 | 1 | 4.0(0) | 4.0 | 4.0 | 24.3(3.1) | 27.0 | 21.0 | 6.3(0.6) | 7.0 | 6.0 |
|  | 2 | 3.0(0) | 3.0 | 3.0 | 24.7(2.1) | 27.0 | 23.0 | 5.0(1.0) | 6.0 | 4.0 |
| 5 | 1 | 3.7(0.6) | 4.0 | 3.0 | 32.0(3.5) | 34.0 | 28.0 | 5.3(0.6) | 6.0 | 5.0 |
|  | 2 | 3.7(0.6) | 4.0 | 3.0 | 25.3(2.5) | 28.0 | 23.0 | 5.7(0.6) | 6.0 | 5.0 |
| 6 | 1 | 3.0(0) | 3.0 | 3.0 | 33.7(2.5) | 36.0 | 31.0 | 5.3(0.6) | 6.0 | 5.0 |
|  | 2 | 3.3 (0.6) | 4.0 | 3.0 | 20.0(2.0) | 22.0 | 18.0 | 6.7(0.6) | 7.0 | 6.0 |

Inspection of the co-membership matrix of the maximal cliques provides some insights into groupings of high and low density within the pens (Figure 2.d). High-density grouping in the co-membership matrix indicates that pairs or groups of animals spent significant amounts of time together in close proximity within a clique, reflecting strong social co-interactions between those animals. Results of the Mantel test, used to compare clique co-membership matrices within the pen, between the early and later growing periods, revealed distinct patterns between pens in both the early and late growing periods (Figure 4). Overall, the correlations in clique co-membership across the 3 days of the early growing period, in the figure, are relatively low, indicating a weak co-membership in clique structures. These results suggest that during the early phase, individual proximity patterns are dynamic, with no consistent grouping trends emerging across days. In contrast, correlations across the 3 days in the later growing period were generally stronger and positive. Furthermore, the correlations between matrices of the early and late growing periods were relatively low, ranging between 0.20 to 0.44 in all pens (except for pen 6, where lower than 0.20 correlation estimates were observed), indicating that cliques were formed of different animals in the early compared to late growing period.

**Figure 4:**
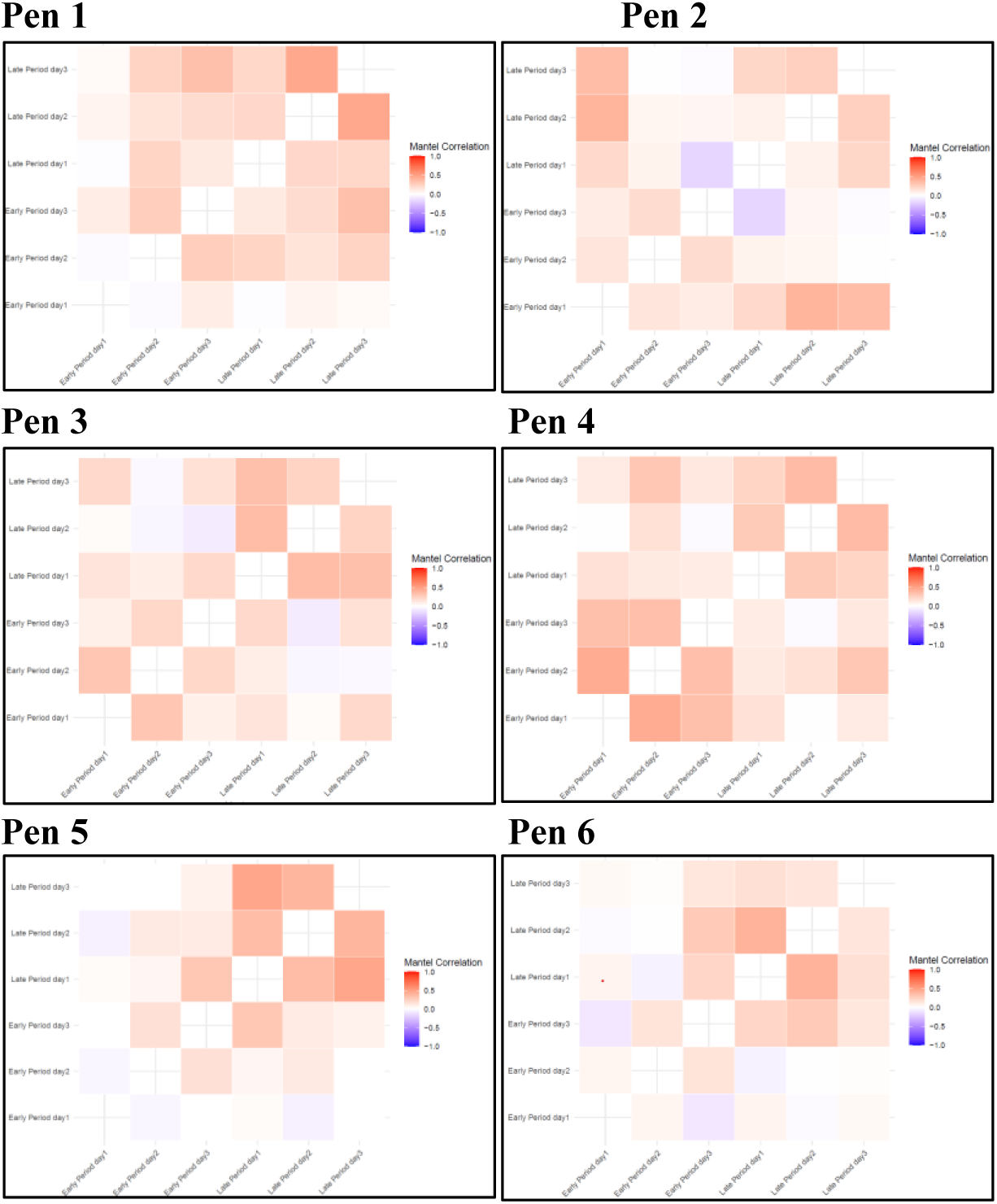
Mantel test for clique co-membership correlation between different days in the early growing period and late growing period for the six studied pens.

### 3.3. Individual SNA traits

Table 4 presents the descriptive statistics of the individual SNA traits across the two growing periods. Overall, the individual SNA traits were stable over the two growing periods, except for closeness centrality and clustering coefficient (Figure 5). Specifically, the mean degree centrality for individuals remained stable across both periods, suggesting that the overall connectivity of individuals did not change over time. Similarly, no significant difference was observed in betweenness centrality between periods.. Although eigenvector centrality showed a slight decrease from the early to the late growing period, this was not statistically significant. On the other hand, closeness centrality significantly increased from the early to the later growing period (p < 0.00001), although closeness was very low at both timepoints. The observed change indicates a notable shift in the proximity network structure across the growing periods, where the direct interactions of the central individuals with other members of the network increased. This shift could be due to the more limited space available in the pen as the pigs matured. Also, the clustering coefficient increased significantly from the early to the late growing period (p = 0.00006), indicating that individuals in the network tended to form more cohesive groups as time progressed.

**Figure 5:**
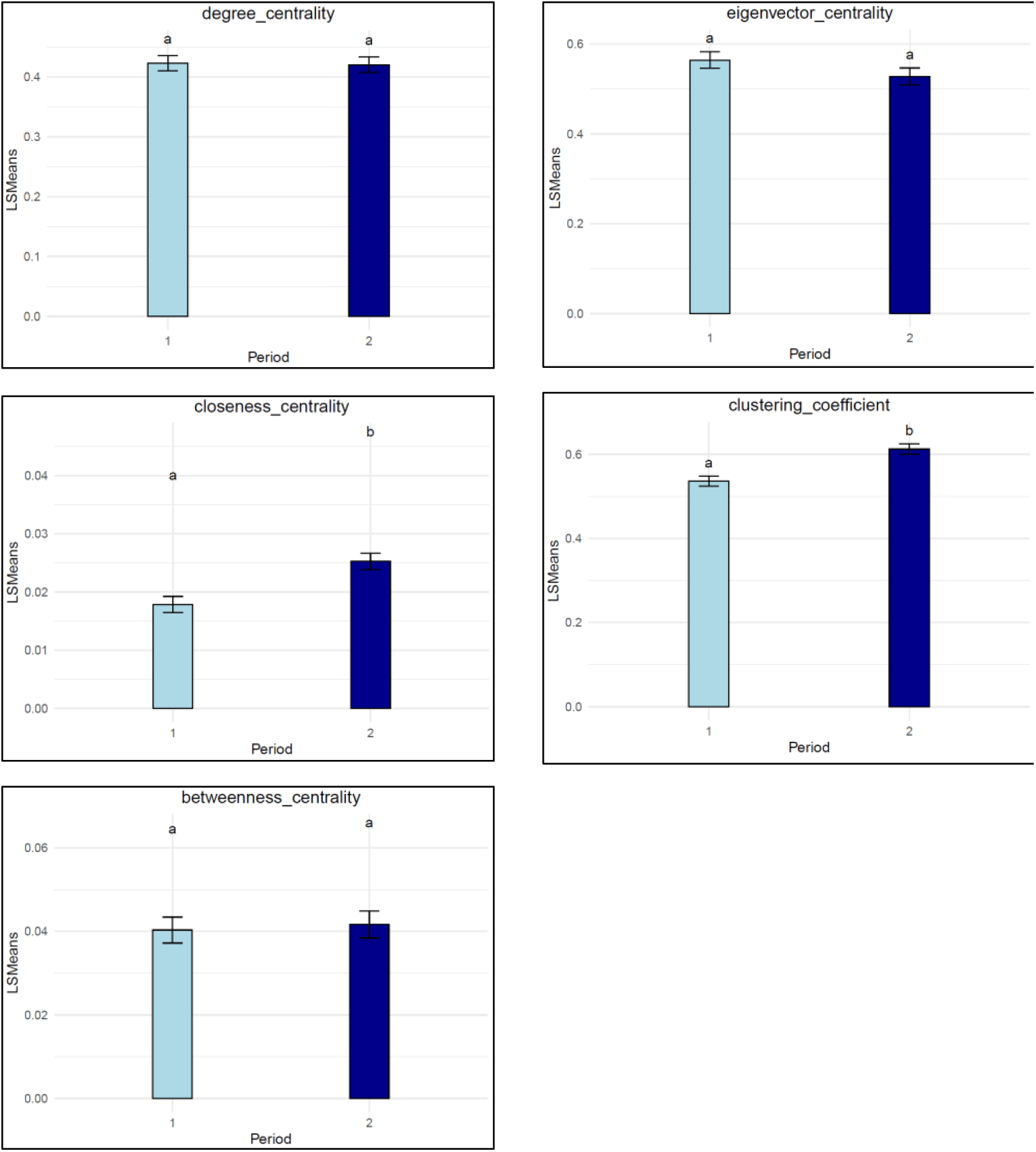
Least square means of the individual SNA traits in different growing periods. Periods not connected by the same letter are significantly different at (P < 0.05).

**Table 4.**
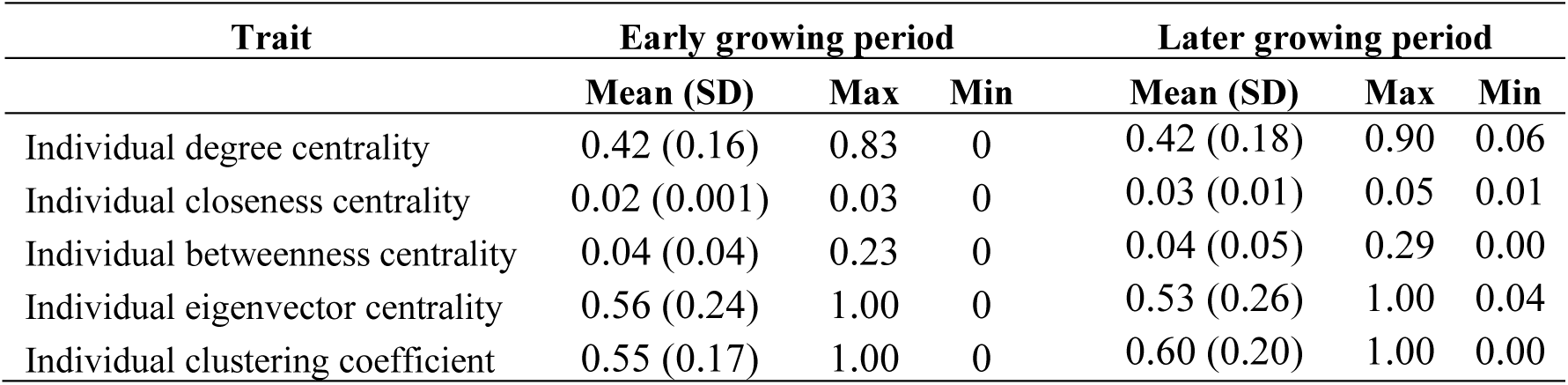
Descriptive statistics for the individual SNA traits of the studied pens for early and late growing periods.

| Trait | Early growing period |  |  | Later growing period |  |  |
| --- | --- | --- | --- | --- | --- | --- |
|  | Mean (SD) | Max | Min | Mean (SD) | Max | Min |
| Individual degree centrality | 0.42 (0.16) | 0.83 | 0 | 0.42 (0.18) | 0.90 | 0.06 |
| Individual closeness centrality | 0.02 (0.001) | 0.03 | 0 | 0.03 (0.01) | 0.05 | 0.01 |
| Individual betweenness centrality | 0.04 (0.04) | 0.23 | 0 | 0.04 (0.05) | 0.29 | 0.00 |
| Individual eigenvector centrality | 0.56 (0.24) | 1.00 | 0 | 0.53 (0.26) | 1.00 | 0.04 |
| Individual clustering coefficient | 0.55 (0.17) | 1.00 | 0 | 0.60 (0.20) | 1.00 | 0.00 |

## 4. Discussion

SNA is widely recognized as a valuable tool for investigating the social structures of animals, including livestock and aquaculture species [43]. Despite its potential, practical applications have often been hampered by insufficient data, as manual data collection is time-consuming and prone to observer bias [8]. However, recent advances in automated monitoring systems have significantly enhanced the ability to capture the position and activities of animals, offering a high degree of reliability and accuracy, even in complex farm environments with factors such as motion blur and varying lighting conditions [22]. The automated system and AI routines that provided the data for the current study were previously subjected to extensive quality control to confirm their reliability in recording and tracking pig position, posture and activities [24,28,29]. However, further validation analyses were also carried out in this study to ensure the reliability of this data (See validation, in the Suppl. Material). The results of this validation showed that the automated data, derived from DL algorithms, demonstrated a matching accuracy for the posture and activity of over 97% with a human observer, with the majority, i.e., 99%, of the errors observed in sitting/lying postures. Similar accuracies, ranging between 95 to 98%, were also reported in previous studies of automated monitoring system used to classify pig posture and activities [22]. Furthermore, to validate the XY coordinates of this data, the size of the animals was simply calculated as the Euclidean distance between the shoulder and rump XY coordinates (Figure S1.a). The repeatability estimates for the animal size, calculated as the within-individual repeatability coefficient of variation, ranged from 0.10 to 0.18 across pens (Table S1). This variability could be due to the change in animal positioning and posture, or measurement errors in the DL system [44,45]. These repeatability estimates suggest that the XY position data are sufficiently accurate for the construction of reliable contact networks based on proximity estimates derived from these position data, with corresponding error margins. Furthermore, the size of the same animal was then compared in different pen locations to evaluate the effect of potential distortion in video camera images on the derived XY coordinates (Figure S1.b). The least square means showed that the difference between the estimates for the inner and outer area of the pen never exceeded 0.15-meter Euclidean distance across all the studied pens (Figure S2). Based on these observed small variabilities during the validations of XY coordinates, proximity was defined in this study as a Euclidean distance of less or equal to 0.5 meters between the shoulders of “standing” pigs, sustained for a duration exceeding the average interaction time between each pair of individuals within each pen each day (Table 1). This definition accounted for measurement variations in the XY coordinates and aligned with previously reported proximity distances (0.3–1 meter) used to define interactions when standing [35,46]. Furthermore, proximity durations exceeding the daily average pen proximity were considered to further address these variations, ensuring that only sustained interactions were included in the construction of the network, while filtering out incidental or brief social proximity.

The aim of this study was to demonstrate the feasibility of constructing informative social contact networks from proximity measures derived from the automated position data. To this purpose, a computational pipeline was developed to show that SNA applied to automated real-time position data facilitates a detailed observation and characterisation of social interactions between pigs within a pen by converting automated data into visual graphs and quantitative network measures. In particular, the study demonstrates the capability of SNA to reveal the strength and stability of interactions, community divisions, and subgroup cohesion within a pen’s social structure (Figure 2).

### 4.1. Group-level SNA

SNA offers the potential of quantifying the social structure of groups, using the social interaction data, providing informative group-level SNA traits [13]. Here, group level SNA showed a notable change in social network dynamics over two growing periods, emphasizing the evolution of social interactions among individuals over time (Figure 3). Group-degree centralization, closeness centralization and betweenness centralization increased significantly between the early and later growing period. The increase in group-level centralization suggests that, over time, fewer animals occupied a highly central position with respect to proximity, whilst more animals adopted a peripheral network position and showed proximity with a restricted number of group members. This may reflect the growing familiarity between animals and the establishment of a clearer social hierarchy within the pen. These results are in accordance with previous studies which have shown that pigs adapt to stable group settings by forming stronger and more consistent social bonds[47]. This adaptation suggests that pigs’ social structures evolve as they become more accustomed to their environment, which could have a direct effect on the expression of harmful behaviour e.g., aggressive interactions.

In contrast, eigenvector centralization and modularity showed no significant change from the early to the later growing period (p = 0.12), and the network density also remained stable across both periods (p = 0.79). This stability suggests that, despite individual changes in social roles, the core structure of proximity behaviours remained consistent throughout the growing periods. This could be attributed to the controlled environment in which the studied pigs were raised, which may have minimized disruptions to their social dynamics. Similar findings have been reported in previous studies, where the lack of significant environmental stressors may have allowed the pigs to maintain their established social patterns over time. However, it would be interesting to assess the consequences on the group-level centralization under changing environments, such as dietary or health challenges or re-mixing of pigs and in particular removal of high centrality individuals from the pen in future investigations.

### 4.2. Communities and cliques

The number of communities remained relatively stable across the early and late growing periods (p = 0.14) (Figure 2.b and 3), as did the size of the largest clique. Maintaining similar community structures over time may indicate a relatively stable social structure within a pen, where pigs continue interacting with the same individuals across both periods. On the other hand, the significant reduction in the number of maximal cliques from the early to the late period highlights a marked shift in the network’s subgroup structure. Maximal cliques represent fully connected individuals, and the decrease in their number during the late growing period suggests that the social network became less concentrated within certain cliques of individuals [11].

Forming cliques has been found to be beneficial for the stability of a group. For instance, [17] observed that pens with cliques, during the 24 hours post mixing, had lower rates of aggression related injuries three weeks later. However, the same authors also reported that the members of such cliques would suffer a higher rate of injuries, compared to the non-clique-members in the first 24 hours. Similarly, at the genetic level, Agha et al. [19], reported that there is a high positive genetic correlation between clique membership in aggression networks and anterior skin lesions 24 hours post mixing, but a strong negative genetic correlation with anterior skin lesions after three weeks post mixing. This highlights the importance of identifying the key animals that frequently participate in such cliques, to increase social stability within a pen. Given this importance, a comprehensive co-membership analysis was conducted for the maximal cliques in the current study. This analysis, along with the visualization of cliques, provided a more detailed insight into the patterns of intense social interactions within the network (Figure 2.d). Specifically, the Mantel test results, comparing clique co-membership matrices between the early and late growing periods, provided a deeper insight into the development of social cliques among pigs through different growth stages. In the early growing period, there was a relatively low correlation between days across all pens, suggesting that pigs exhibit less defined social relationships, especially when forming cliques. The dynamic nature of these early relationships likely reflects the exploratory behaviour of pigs during the initial stages of group formation when dominance hierarchies and stable social groups have yet to be fully established [48,49]. Higher correlations during the later growth stages and the low correlations observed in clique co-membership matrices between the two growth periods imply that pigs tend to maintain more stable proximity patterns over time (Figure 4). This trend toward stability in social grouping aligns with research showing that pigs form more cohesive social groups over time, as they establish a relatively stable dominance hierarchy and potentially preferential affiliative relationships become more established [50]. The increasing stability in clique structures could reduce social tension and aggression, which is advantageous for welfare and productivity as pigs approach market weight [51]. Understanding these patterns is important for improving group housing systems in commercial pig production.

It is also noteworthy that the co-membership analyses revealed differences in social patterns across groups. In this study pen 6 was an exception to the above, where the Mantel correlations remained lower compared to other pens, suggesting that the pigs in this pen may have experienced different social dynamics. These variations could be attributed to factors such as changes in environmental or physical conditions (e.g. disease), or to the mix of individual personalities and attributes such as cognitive ability [52]. Further investigation into the specific conditions could help explain these discrepancies.

Finally, it should be noted that proximity when active could indicate engagement in positive (affiliative) social or in negative social interactions (such as aggression). Under commercial farming conditions, the formation of communities and cliques in the pen could be partly influenced by the distribution of feeding and water resources, group size and pen type. Although this was not explicitly considered in the current study, this hypothesis could be further investigated by constructing social networks for the area around these resources and performing community detection and clique analyses. By identifying the high- and low- density areas associated with resource locations, adjustments to the placement of resources could be made in order to reduce competition and harmful behaviours among pigs.

### 4.3. Individual SNA traits

SNA also offers informative individual measures that quantify the role of individuals in social interactions [10]. The population-level stability of key individual SNA traits such as degree centrality and betweenness centrality over the two growing periods. Stability in degree centrality implies that socially influential individuals maintain their status over time, which can be leveraged for management and breeding purposes. Likewise, the consistency in betweenness centrality indicates that key pigs consistently bridge different parts of the social network over the growing period. This stability could be useful for group management when identifying pigs that play critical roles in the social structure, ensuring that these animals are strategically managed to maintain network cohesion. This stability also implies that disease transmission within a pen may persistently operate through the same key individuals [53].

On the other hand, the significant increase in individuals’ closeness centrality and clustering coefficient observed from the early to late growing periods suggests that as pigs mature, their social dynamics evolve. This change may be linked to the pigs’ adaptation to their environment and the establishment of more defined social relationships as they grow. Similarly, the increase in clustering coefficient reflects the formation of more cohesive subgroups over time, likely as pigs become more familiar with their pen mates and develop stronger social bonds. The significant changes in these traits indicate that proximity behaviours shift as pigs adapt and mature. Similar results were also observed in primates [54] and wild animals [55].

#### Implications for management and breeding

This proof-of-concept study has demonstrated that integrating AI assisted automated monitoring technologies with SNA provides an efficient, real-time method for analysing animal social interactions, advancing beyond traditional manual observation techniques. While this study focused on pigs, utilizing our data and experience with this species, the methods developed can be broadly applicable to other livestock and aquaculture species. The decreasing cost of monitoring technologies and the growing adoption of automated systems in these industries yield valuable data that requires innovative analytical approaches for deriving quantitative measures that can be effectively used to improve the management and/or breeding of farm animals [9,10]. This information could potentially be used for optimizing group compositions to reduce competition or increase positive interactions between animals in the pen. Furthermore, since individual SNA traits are heritable and favourably correlated with key economic traits, incorporating them into breeding programs could potentially improve the accuracy of predicting social compatibility of animals in commercial farms [16,19].

In the welfare context, combining the application of SNA and automated monitoring extends beyond productivity to potentially significantly enhance animal well-being in commercial farms [15]. Understanding the social dynamics within a pen allows for the early identification of harmful social behaviour e.g., aggression or tail biting [56]. Also, the community detection and identification of clique members can be used at the farm level to create more homogeneous groups, minimize stress and promote positive interaction between animals. Improved welfare not only addresses ethical responsibilities but also has a direct impact on the long-term productivity of farm animals [6].

In health and disease management, automated monitoring and sensor technologies have emerged as a promising tool for tracking and managing disease incidences in farm animals [57]. Integrating SNA with automated data thus offers a powerful tool for understanding disease transmission and managing it in farm animals [53]. For instance, identifying key individuals, e.g. those with high degree and betweenness centrality, could help map transmission pathways. These traits also enable early and targeted interventions, such as detecting animals exhibiting abnormal interactions, such as withdrawal from social contact that might indicate sickness behaviour, and adjusting group compositions to limit disease spread [58]. This proactive approach to health management not only reduces disease transmission but would also help in reducing the excessive use of antibiotics and vaccines, contributing to more sustainable farming practices. Scientifically, SNA could be used to parameterise epidemiological models of infection spread, allowing for more accurate model predictions [53].

However, it is important to note that this study focused on non-directional weighted networks based on the duration of proximity between pigs in a standing position. While such proximity data can be significant for disease transmission studies [59], they may not fully capture the complexity of social interactions related to productivity and welfare issues. Therefore, to gain a more comprehensive understanding, directed network such as those reflecting feeding interactions or aggressive behaviours, should be investigated in future studies, as they may provide more insights into social dynamics [15]. By breeding pigs for desirable social traits, producers can enhance group cohesion and overall health, leading to more efficient production systems and improved animal welfare.

## Conclusion

This proof-of-concept study demonstrates the feasibility and potential of integrating SNA with automated monitoring data and AI tools to construct informative social contact networks. By evaluating group-level SNA traits, identifying community structures, analysing clique co-membership, and assessing individual SNA traits, we gained novel insights into the social interactions of group-housed pigs under commercial farm conditions, and how these change over time. These novel insights highlight the promise that SNA offers for optimizing management practices and improving animal performance, health, and welfare of farm animals.

## Institutional Review Board Statement

**Conflicts of Interest:** The authors declare no conflict of interest.

