## Supplementary material for "Revealing the hidden social structure of pigs with AI assisted automated monitoring data and social network analysis": Suppl.

***Validation of the automated data***

To validate the system's accuracy in determining individual animals' location, posture, and activity, 36 pigs, six from each of the six pens, were selected at random and marked with distinct colours for easy recognition from images. Over the following five days, the tracking algorithm produced ~50 annotated images per pen, capturing each pig's ear tag identification, posture, activity and position. Posture (lying, sitting, standing), activity (eating and drinking) and position (XY coordinates of shoulder and rump), and data for the 36 marked pigs were then extracted from the DL algorithm and manually cross-referenced with the corresponding images. Specifically, as initial validation of the XY coordinates derived from the DL algorithms, animals were sorted based on their X and Y shoulder and rump coordinates, and the relative position of animals to each other within the data files was compared with their relative positions within the images (Figure S1). The results of this validation showed that the automated data, derived from DL algorithms, demonstrated a matching accuracy for the posture and activity of over 97% with a human observer, with the majority, i.e., 99% of the errors observed in sitting/lying postures (Data not shown). Notably, the validation of the relative XY position data, revealed no errors in the derived data, further affirming the robustness of the DL methods.

As the social networks constructed in this study are based on individuals' proximity to each other, calculated from the position data as outlined in the next section, XY shoulder and rump coordinates were further validated for their precision, reliability and potential bias. This was done by deriving an estimate of each animal's size from these coordinates, and by assessing the stability of the size of each animal in the six pens across space and within 3-day periods, the early growing period (the first month after mixing) and a later growing period (60 days after mixing), where one would expect the size to remain constant within each of these time periods. To this purpose, the size of the animals was simply calculated as the Euclidean distance between the shoulder and rump XY coordinates (Figure S1.a). The descriptive statistics for the size of the animals, calculated based on this Euclidean distance, for each pen within the studied 3-day period in the first month and 3 days, 60 days later is shown in Table S1. The repeatability estimates for animal size, calculated as the within-individual repeatability coefficient of variation of animal size, ranged from 0.10 to 0.18 across pens. A smaller value Repeatability CV indicates higher repeatability. These moderate values suggest consistency in measurements, but small fluctuations likely arose from animal positioning, posture changes, or measurement errors in the DL system. These repeatability estimates suggest that the XY position data are sufficiently accurate for the construction of reliable contact networks based on proximity estimates derived from these position data, with corresponding error margins.

Furthermore, to evaluate the effect of potential distortion in video camera images on the derived XY coordinates, the pen was divided into two distinct areas: the inner and outer area of the pen, where the inner area was defined by X-coordinates ranging from 1.5m to 4.5m and Y-coordinates ranging from 1m to 2m, measured from the top-left corner of the pen (Figure S1.b). The size of the same animal was then compared when located inside versus outside this defined area using mixed linear models with the designated area as fixed effect, fitting pen and animal as random effects. While the mixed model analysis revealed statistically significant differences between the size estimates associated with the inner and outer area of the pen ($p < 0.0001$ for all comparisons), the least square means showed that the difference between the estimates for the inner and outer area of the pen never exceeded 0.15-meter Euclidean distance across all the studied pens (Figure S2). This difference was taken into consideration when defining proximity thresholds used in the social network analysis in the subsequent steps, as proximity between animals within the same contemporary group was defined by a threshold of 0.5 meters Euclidean distance between the shoulders, sustained for a duration longer than average interaction time between individuals within each pen each day (Table S2). These chosen thresholds accommodate for potential measurement errors in the XY coordinates. Furthermore, the chosen 0.5 meters distance is within the range of previously identified proximity distances between 0.3 to 1 meter defining interactions during standing position [34].

**Table S1. Descriptive statistics for the size of the animals calculated based on the Euclidean distance between shoulder and rump XY coordinates for each pen per growing period.**

| Growing Period | Pen | No. of animals | Mean | SD | Min | Max | Repeatability CV |
|---|---|---|---|---|---|---|---|
| **Early** | 1 | 19 | 0.49 | 0.05 | 0.00 | 0.89 | 0.11 |
| | 2 | 19 | 0.48 | 0.05 | 0.00 | 0.79 | 0.11 |
| | 3 | 18 | 0.45 | 0.05 | 0.00 | 0.87 | 0.11 |
| | 4 | 19 | 0.44 | 0.04 | 0.00 | 0.81 | 0.10 |
| | 5 | 19 | 0.49 | 0.05 | 0.00 | 0.83 | 0.10 |
| | 6 | 19 | 0.47 | 0.05 | 0.00 | 0.74 | 0.10 |
| **Late** | 1 | 19 | 0.73 | 0.10 | 0 | 1.24 | 0.14 |
| | 2 | 18 | 0.70 | 0.11 | 0 | 1.06 | 0.15 |
| | 3 | 16 | 0.71 | 0.09 | 0 | 1.23 | 0.13 |
| | 4 | 17 | 0.70 | 0.09 | 0 | 1.18 | 0.12 |
| | 5 | 19 | 0.72 | 0.13 | 0 | 1.12 | 0.18 |
| | 6 | 18 | 0.73 | 0.09 | 0 | 1.13 | 0.12 |

**Table S2. Descriptive statistics of the proximity time between each pair of animals in each day in the two growing periods (Early=1 and late=2).**

| Growing Period | Pen | Day | No of animals | Proximity time | | |
|---|---|---|---|---|---|---|
| | | | | Mean | Max | Min |
| **Early** | 1 | 1 | 19 | 34.04 | 117.26 | 0.90 |
| | | 2 | 19 | 28.96 | 93.15 | 0.81 |
| | | 3 | 19 | 30.03 | 92.91 | 0.90 |
| | 2 | 1 | 19 | 28.04 | 98.81 | 2.12 |
| | | 2 | 19 | 31.97 | 85.32 | 2.59 |
| | | 3 | 19 | 19.37 | 70.77 | 0 |
| | 3 | 1 | 18 | 39.13 | 124.1 | 3.7 |
| | | 2 | 18 | 25.77 | 76.79 | 2.49 |
| | | 3 | 18 | 26.89 | 82.52 | 3.26 |
| | 4 | 1 | 19 | 28.68 | 90.74 | 1.08 |
| | | 2 | 19 | 21.44 | 83.04 | 0 |
| | | 3 | 19 | 23.61 | 85.34 | 0 |
| | 5 | 1 | 19 | 27.01 | 120.28 | 0.16 |
| | | 2 | 19 | 31.04 | 97.46 | 3.15 |
| | | 3 | 19 | 19.29 | 94.5 | 0 |
| | 6 | 1 | 19 | 27.13 | 82.91 | 1.34 |
| | | 2 | 19 | 18.62 | 61.77 | 1.06 |
| | | 3 | 19 | 19.59 | 59.49 | 0 |
| **Late** | 1 | 1 | 19 | 17.12 | 63.24 | 0.27 |
| | | 2 | 19 | 29.72 | 98.15 | 3.24 |
| | | 3 | 19 | 25.94 | 76.93 | 1.13 |
| | 2 | 1 | 18 | 24.68 | 87.82 | 2.71 |
| | | 2 | 18 | 18.92 | 62.62 | 0.19 |
| | | 3 | 18 | 16.37 | 47 | 3.47 |
| | 3 | 1 | 16 | 17.63 | 57.61 | 0 |
| | | 2 | 16 | 19.86 | 67.47 | 0.52 |
| | | 3 | 16 | 8.75 | 56.04 | 0 |
| | 4 | 1 | 17 | 24.06 | 84.12 | 0.62 |
| | | 2 | 17 | 23.32 | 82.47 | 0.91 |
| | | 3 | 17 | 28.86 | 95.58 | 4.96 |
| | 5 | 1 | 18 | 15.99 | 75.07 | 0 |
| | | 2 | 18 | 19.02 | 71.38 | 0 |
| | | 3 | 18 | 11.5 | 38.06 | 0 |
| | 6 | 1 | 18 | 16.57 | 50.82 | 0.25 |
| | | 2 | 18 | 21.26 | 127.07 | 1.17 |
| | | 3 | 18 | 12.16 | 47.84 | 0 |

**(a)**

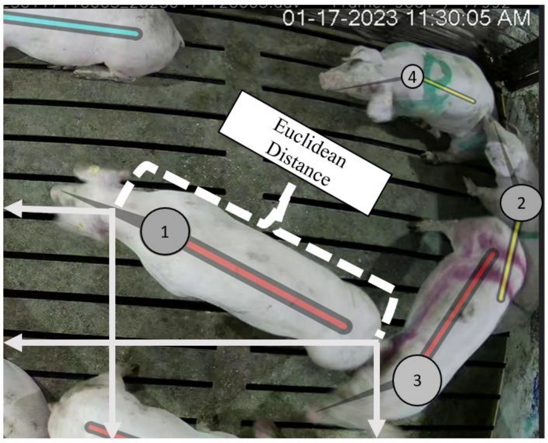


**(b)**

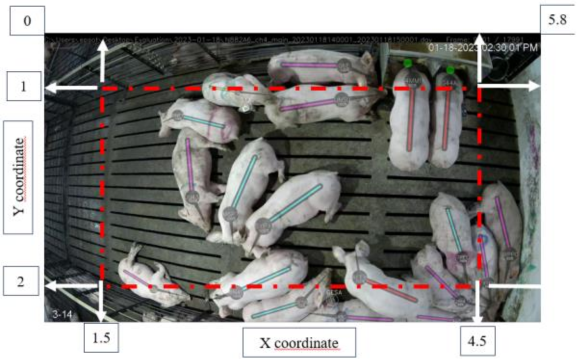


**Figure S1. (a)** The calculation of the animal size based on the Euclidean distance calculated from the XY coordinates of the shoulder and rump, **(b)** Validation for potential distortion in video camera images by dividing the pen into two distinct areas based on XY coordinates. The inner area of the pen is defined by X-coordinates ranging from 1.5 m to 4.5 m and Y-coordinates ranging from 1 m to 2 m, measured from the top-left corner of the pen.

### (a) Early growing period

#### Pen 1

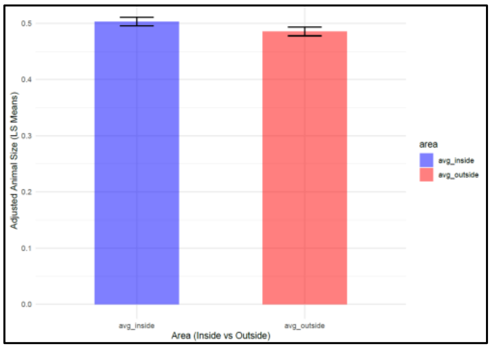


#### Pen 2

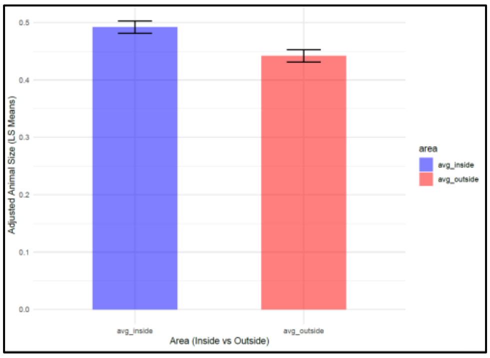


#### Pen 3

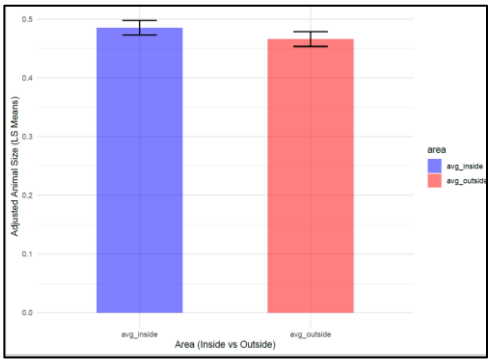


#### Pen 4

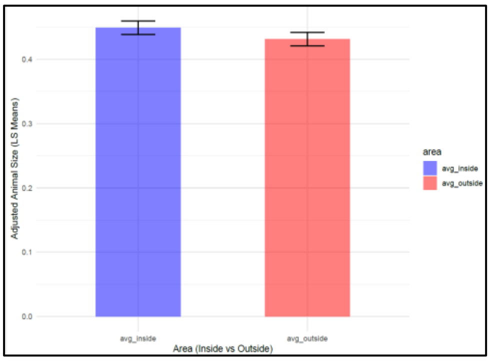


#### Pen 5

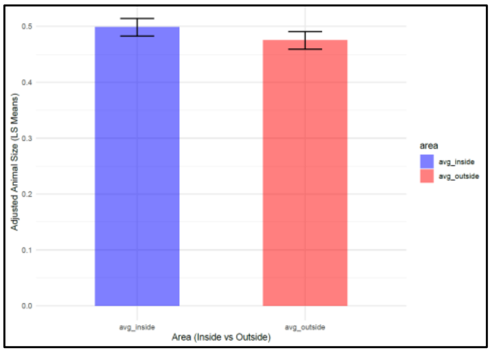


#### Pen 6

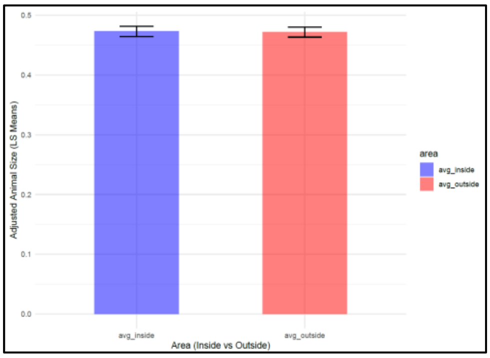


### (b) Late growing period

#### Pen 1

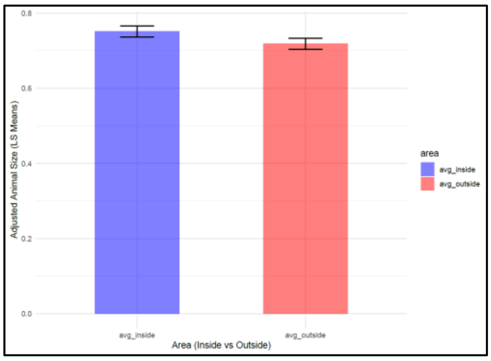


#### Pen 2

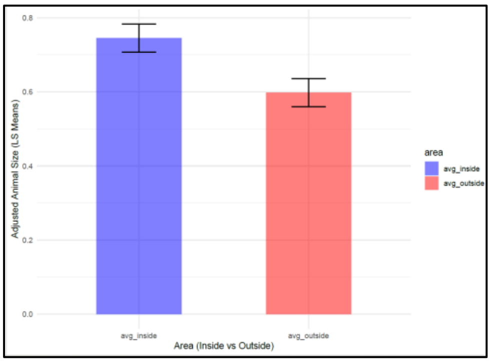

**Pen 3**

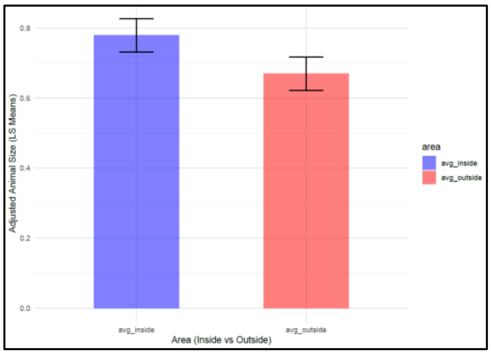


**Pen 4**

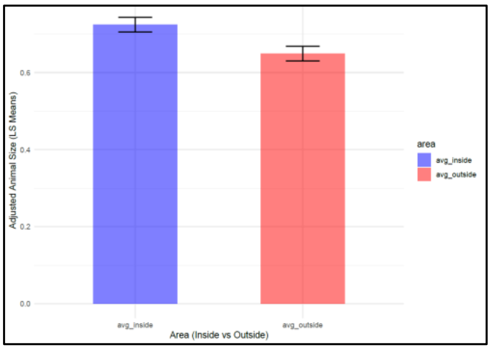


**Pen 5**

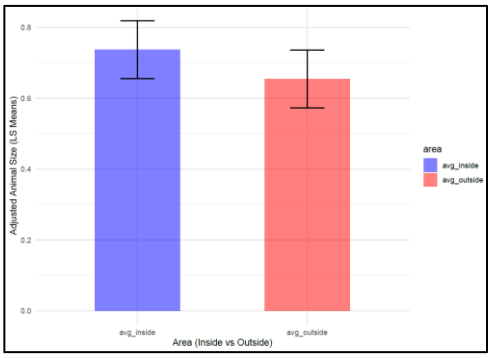


**Pen 6**

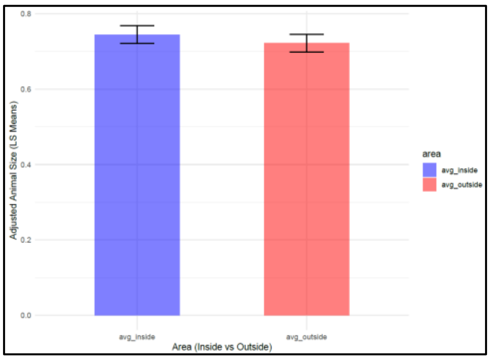


**Figure S2. Least square means from the mixed model analysis of size estimates corresponding to the inner (in blue) and outer areas (in red) of the pen for (a) the early and (b) late growing periods.**
